# Coronary atherosclerotic plaque composition and classification in hypercholesterimic pigs

**DOI:** 10.1101/2025.06.06.658319

**Authors:** Mingqiao Li, Patrice Delafontaine, Sergiy Sukhanov

**Affiliations:** Tulane University School of Medicine, Department of Medicine, New Orleans LA 70112

**Keywords:** coronary atherosclerotic plaque, plaque classification, plaque morphology, plaque vulnerability, large animal model of atherosclerosis, atherosclerotic pigs, Rapacz pigs

## Abstract

Rapacz pigs with familial hypercholesterolemia (FH pigs) fed with high-fat diet (HFD) develop early atherosclerotic lesions and complex atheromas in coronaries mimicking human coronary atherosclerotic disease (CAD). FH pigs have proven to be an excellent model for basic and pre-clinical atherosclerosis-focused research. However, unlike human atherosclerosis there has been no established classification system for porcine atherosclerosis. We isolated 104 plaque-containing coronary fragments from atherosclerotic FH pigs. A set of indices (features) of vessel and plaque morphology were quantified for each plaque, including intima-media ratio, vessel size, necrotic core area and fibrous cap thickness. Multifeatured clustering algorithm identified 4 clearly distinguishable plaque groups (A-D). Plaque cellular composition was assessed by immunohistochemistry to quantify relative level of smooth muscle-like, endothelial-like and macrophage-like cells. Plaque neovascularization, collagen levels, calcification and features of vulnerable plaque were assessed and used as additional numerical criteria for plaque classification and to establish the similarity of porcine plaque to specific types of human lesions. Our results indicate that porcine plaque type A, B, C and D correspond to human type III (intermediate lesion), type IV (atheroma), type V (fibroatheroma) and type VI (high-risk vulnerable plaque), respectively. Overall, our data demonstrates the suitability of using the FH pig as a pre-clinical model of human-like coronary atherosclerosis with great potential to advance emergent research in the field of CAD, especially in study of vulnerable plaque and in discovery research.

## INTRODUCTION

Atherosclerosis is a chronic inflammatory disease in which there is buildup of plaques inside coronaries ranging from early lesions to advanced atheromas^1^. Atherosclerosis is characterized by the accumulation of lipids, fibrous elements, and calcification within the coronary arteries. This process is initiated by endothelial activation, followed by formation of early lesions, activation of inflammatory pathways leading to fibrous plaque formation, and development of complex-appearing lesions containing large necrotic cores and fibrous caps. Coronary plaque progression involves degradation of vascular endothelial cells (EC) compromising intima integrity, depletion of smooth muscle cells (SMC) and an increased number of macrophages (MF). Advanced atheroma development is also associated with marked change in the vessel morphology, including an increase in coronary size (outward remodeling, compensatory response to atherosclerotic luminal narrowing), and thinning of vascular media. Ultimately, coronary atherosclerotic disease (CAD) can result in increased plaque vulnerability, plaque rupture, exposing thrombogenic components of the plaque, activating the clotting cascade and promoting thrombus formation. Alternatively, atherosclerotic disease initiates plaque erosion, which is characterized by an intact fibrous cap and the presence of a platelet-rich thrombus and neutrophil infiltrates. Altogether, these processes result in cardiovascular complications that remain the leading cause of death worldwide.

To develop a standard framework of histopathological morphologies of lesions for investigating the mechanisms of the disease, classifications of human coronary lesions were proposed^2–5^. The Committee on vascular lesions of the Council on Arteriosclerosis, American Heart Association (AHA) developed a classification of human atherosclerotic lesions based on the histological and histochemical composition^5, 6^. The classification consists of six categories to include early lesions (type I), adaptive intimal thickening (type II), transitional or intermediate lesions (type III) and advanced plaques characterized as atheroma (type IV), fibroatheroma (type V) and complicated plaques with surface defects, hematoma-hemorrhage, and/or thrombosis (type VI). The plaque classification provided a set of specific morphological criteria to evaluate the effectiveness of drug or other interventions and also to provide a reference to test the suitability of animal models as surrogates for the human disease.

The mouse model has become the predominant one to study experimental atherosclerosis, however the extent to which the mouse serves as an accurate model of the human disease is still debatable^7^. Most murine models do not develop unstable atherosclerotic plaques, and murine plaques have a reduced number of SMC and smaller medial *vasa vasora* compared to humans. Unlike humans, mice do not develop significant atherosclerosis in the coronary arteries but manifest advanced atherosclerotic plaque in the aortic root. Furthermore, mice do not exhibit the same range of HDL subsets as is found in humans^7^. The translation of the knowledge obtained from studies in mice to the development of drugs for human atherosclerosis includes use of porcine models of atherosclerosis. Pigs spontaneously develop atherosclerosis, and this is accelerated by feeding an atherogenic high-fat diet (HFD). The pig has a human-like lipoprotein profile, and develops lesions in the coronary arteries, resembling human lesions in complexity, with a thick fibrous cap, a large necrotic/lipid core, intraplaque neovascularization (IPN), calcification and hemorrhaging^8^. Pigs are the FDA-preferred species for testing human cardiovascular devices and the primary choice for preclinical toxicological testing of anti-atherosclerotic drugs, including statins^9^. Mild atherosclerotic lesions first appear in coronary arteries, and both plaque distribution and composition follow a pattern comparable to that of humans, with early lesions transitioning to complex plaques. Familial hypercholesterolemia is a human genetic disorder with high cholesterol and low-density lipoprotein (LDL) levels, resulting in excessive atherosclerosis^10^. Pigs with familial hypercholesterolemia (FH pigs) have been described and initially characterized by Jan Rapacz^11, 12^. FH pigs have a point mutation in both LDL receptor alleles that reduces receptor binding, and allele variations in apolipoprotein B further contribute to the phenotype. FH swine have been shown to be an excellent model for translational atherosclerosis-related research^8^.

We have demonstrated recently that HFD-fed female FH pigs develop complex lesions in coronaries with features of advanced atheromas^13^ similar to what is reported for patients with CAD. Considering the great potential of FH pigs to provide insight into mechanisms of atherosclerosis and to test novel anti-atherogenic therapies, it is important to formally classify porcine coronary atherosclerosis and its similarity to human disease. To our knowledge, currently there is no coronary lesion classification developed for animal models. Here, we propose the classification of coronary plaques for FH pigs. We used a multi-featured clustering algorithm to identify plaque groups with clearly distinguished phenotypes and assigned these groups to the human lesion classification. In summary, our results provide a first-of-kind coronary lesion classification developed for animal models. Our data establishes the suitability of FH pigs as a pre-clinical model closely mimicking human CAD and provides researchers with an instrument to assess changes in lesion phenotype induced by interventions.

## DATA AVAILABILITY

Please see the Supplemental Material for methods. Data supporting the findings of this study are available from the corresponding author upon reasonable request.

## RESULTS

### Coronary plaque morphometry and plaque clustering

The proximal part of the porcine RCA and LAD was cut into 6 fragments of 5.0 mm length each to generate a set of vascular specimens. A total of 104 coronary fragments were used to quantify indices of vessel and plaque morphology: total vessel cross-sectional area (CSA), tunica media and tunica intima CSA, thickness of tunica media and plaque’s fibrous cap, necrotic/lipid core area. Tunica media and intima CSA were used to calculate intima-media ratio (IMR), surrogate marker of atherosclerosis.

To develop the algorithm for coronary plaque classification sections were sorted by IMR from the lowest (IMR=0.48) to the highest (IMR=22.87). The increase in IMR correlated with visual alteration in plaque phenotype from early lesion to advanced atheroma suggesting that IMR is a potentially feasible criterion for lesion grading. Plaques with IMR in the 1^st^ quartile have neither a fibrous cap (FC) nor necrotic core (NC) and the vessel contained a relatively thick tunica media. Increase in IMR was associated with appearance of NC and FC, followed by gradual elevation in NC area, and marked thinning of vascular media. CAD progression induces outward vessel remodeling (increase in vessel size) associated with the presence of a large lipid/necrotic core, and thin FC, both of which are histological markers for plaque vulnerability^14^. We observed a marked increase in vessel CSA for porcine coronary fragments containing plaque with IMR in the 3rd-4^th^ quartile.

These observations prompted us to perform plaque classification using multiple morphological criteria (features). To discriminate plaque types in porcine coronaries, we chose four criteria: IMR, vessel CSA (mm^2^), NC CSA (mm^2^), and FC thickness (mm). Prior to clustering, the value of each feature was standardized using z-score normalization to ensure equal weighting in the analysis. The 4-featured dataset was submitted for unsupervised K-means clustering to identify plaque groups. The Elbow and gap statistics method showed that the optimal number of clusters in the multifeatured dataset was 4 (Suppl.Fig.2). The clustering algorithm identified the following clusters: A (29 fragments), B (31), C (29) and D (15). Results of clustering analysis, morphometry of coronaries and representative images are shown in Fig.1 and Table 1. Each cluster was clearly distinguished from others by the significant difference in at least 2 features. Specifically, group A was distinguished from other groups by the absence of FC and NC. Group C and D were distinguished from group A and B by increases in both NC area and in the vessel size. In addition, each plaque group has an IMR significantly different from others. To reduce the multi-featured data set to 2-dimensions we used UMAP technique. UMAP confirmed a clear visual pattern of data clustering for all groups (Suppl. Fig.3).

**Figure 1.**
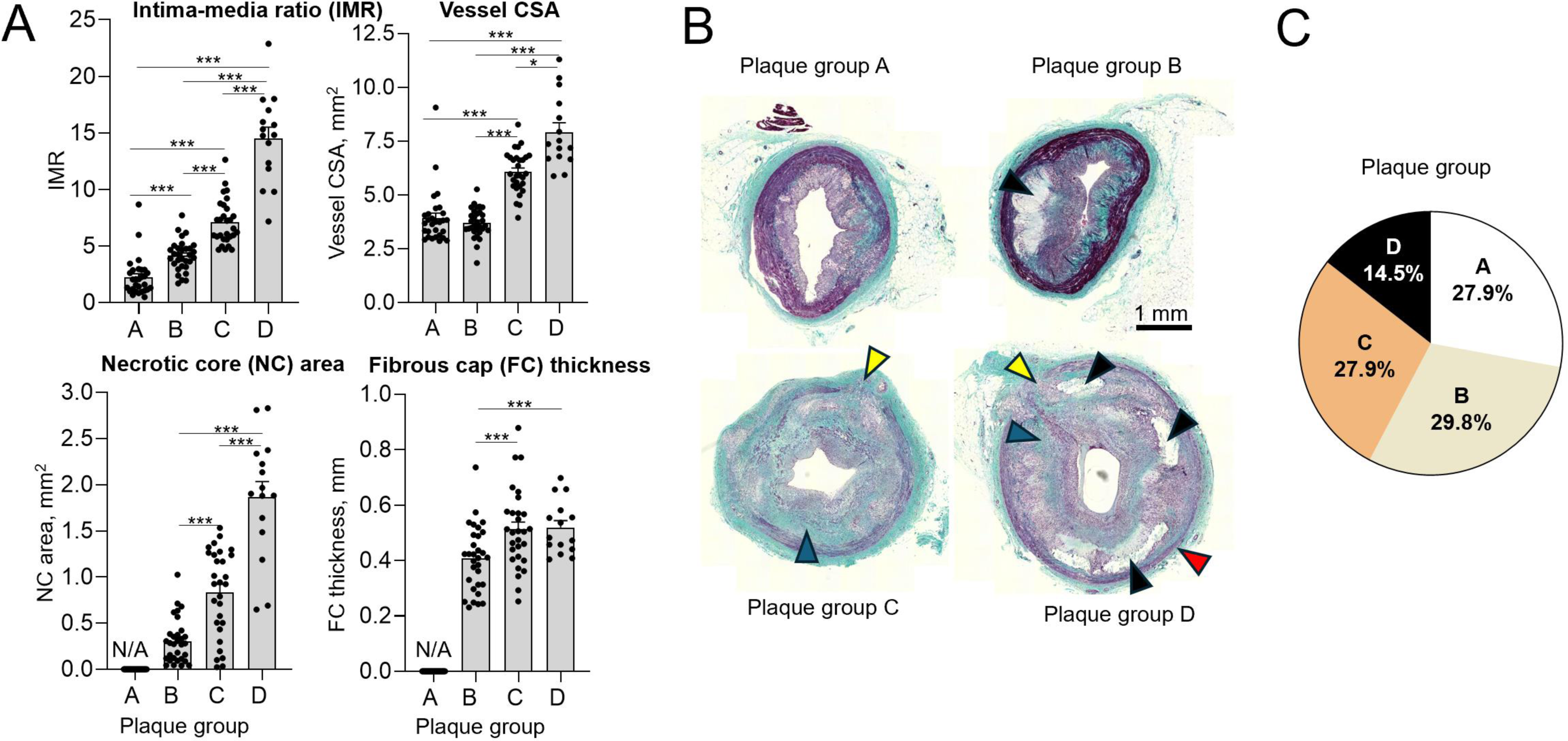
Porcine coronary plaque clustering. Coronary fragments (N=104) were isolated from RCA and LAD of FH pigs, cross-sections were cut and stained with Trichrome stain. The four indices (features) of vessel and plaque morphology were quantified: intima-media ratio (IMR), vessel area (mm^2^), plaque’s fibrous cap thickness (mm) and necrotic core area (mm^2^) and used to identify plaque groups with multifeatured clustering algorithm. The algorithm identified 4 clusters/plaque groups (A-D). A, clustering results are shown for each feature. *P<0.05, ***P<0.005, <u>ttest.</u> B, representative coronary sections. C, Plaque groups as part of whole. Yellow arrows, media breaks; blue arrows, high collagen; red arrow, thin vascular tunica media; black arrow, large necrotic cores. Scale bar, 1mm.

**Table 1.** Coronary arteries morphometry.

| **Plaque group** | **A** | **B** | **C** | **D** |
| --- | --- | --- | --- | --- |
| Intima-media ratio (IMR) | 2.24±0.32 | 4.15±0.26 | 7.09±0.37 | 14.49±1.01 |
| Intima-media thickness, mm | 1.35±0.03 | 1.42±0.02 | 1.79±0.03 | 1.95±0.05 |
| Vessel CSA, mm<sup>2</sup> | 3.91±0.23 | 3.70±0.13 | 6.07±0.18 | 7.92±0.43 |
| Tunica media CSA, mm<sup>2</sup> | 1.32±0.09 | 0.96±0.05 | 1.11±0.07 | 0.85±0.09 |
| Tunica intima CSA, mm<sup>2</sup> | 1.67±0.15 | 2.41±0.11 | 4.62±0.15 | 6.58±0.34 |
| Necrotic core CSA, % | N/A | 12.09±1.57 | 18.81±2.11 | 29.99±3.13 |
| Tunica media thickness, um | 211.5±7.90 | 167.4±6.16 | 156.3±8.82 | 125.5±11.6 |
| Fibrous cap thickness, um | N/A | 406.4±22.2 | 512.6±27.1 | 519.0±24.7 |

### Plaque cellular composition

SMC, EC and MF levels were quantified by immunohistochemistry (IHC) using cell marker-specific antibodies: α-smooth muscle actin (αSMA), CD31 and scavenger receptor A (SRA), respectively. Phenotype switching in atherosclerosis^15^ complicates marker-based cell identification. The IHC protocol was validated by staining of serial porcine sections containing advanced plaque with a set of cell marker antibodies: 4 marker antibodies to detect SMC, 3 antibodies for MF and 3 antibodies for EC. We reported that each SMC marker antibody stained virtually identical cell populations in the plaque or tunica media, and a similar conclusion was made for MF and EC marker antibodies^13^. The IHC protocol includes staining of coronary cross-sections with aSMA, CD31 and SRA antibody allowing the demonstration that cells immunopositive for SRA, were immunonegative for non-MF markers (αSMA, and CD31) and *vice versa* (see Fig.2A and^13^). Overall, these data indicate that antibodies used for the IHC are specific and have virtually no cross-reactivity.

**Figure 2.**
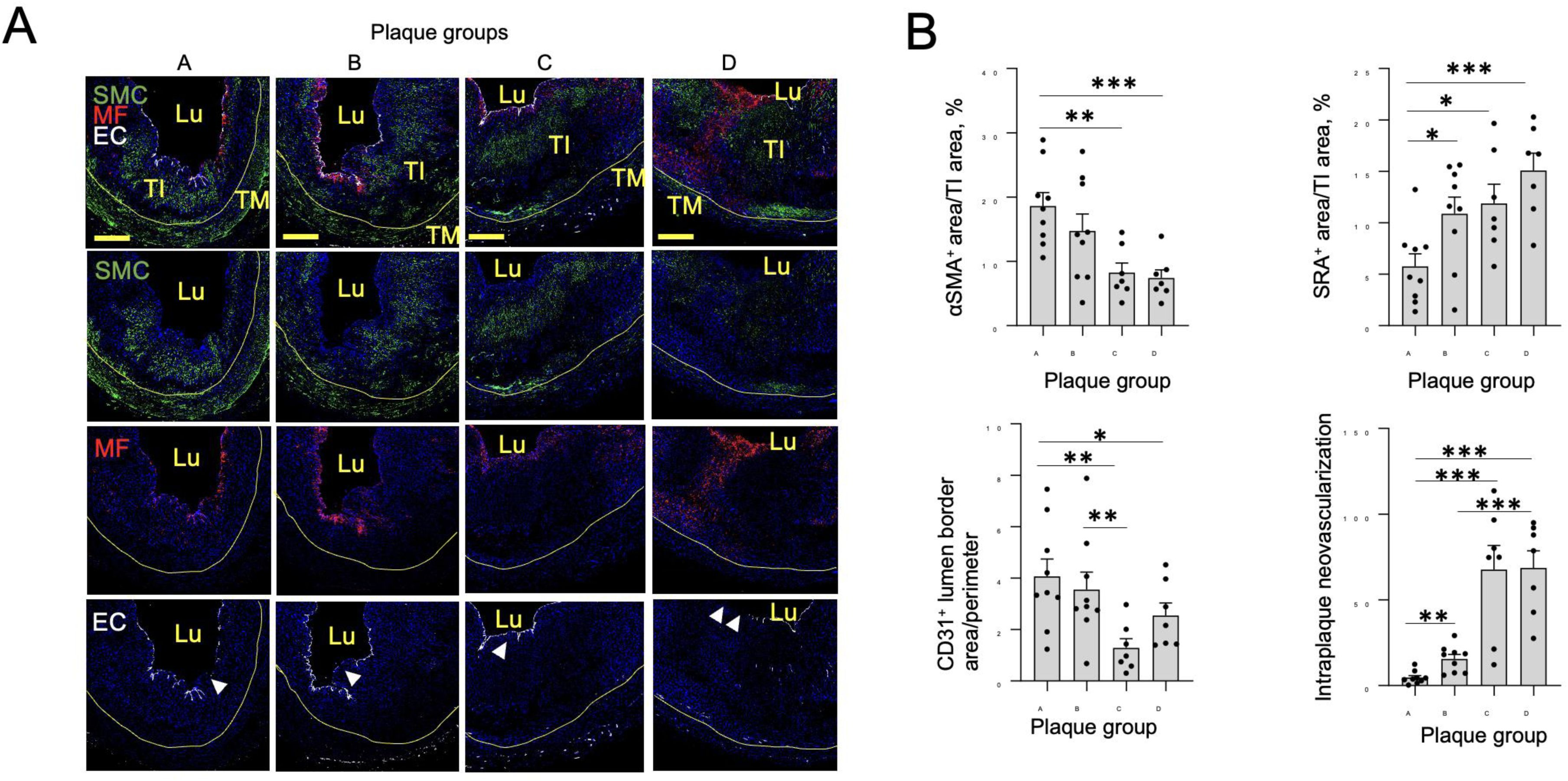
Coronary plaque cellular composition. The coronary sections (N=9/group A; N=9/group B, N=7/group C, N=7/group D) were immunostained with cell marker antibodies (CD31, endothelial cells, EC; scavenger receptor A, SRA, macrophages, MF; a-smooth muscle actin, aSMA, smooth muscle cells, SMC) and DAPI. Primary antibody signal was amplified using fluorescent conjugates. The greyscale images were generated for each fluorescent channel, tunica intima (Tl) was outlined, and area immunopositive for cellular marker was quantified within Tl and normalized per Tl area. lntraplaque neovascularization (IPN) was calculated as CD31+-positivity per plaque after subtracting signal in the luminal border. A, representative images of cell marker-stained sections, B, quantitative data. Yellow scale bar, 500 um. Lu, lumen, TM, tunica media. Yellow curve shows internal elastic membrane, the borderline between TM and Tl. White arrows, breaks in EC layer. *P<0.05, **P<0.01, ***P<0.005, ttest.

Vascular SMC are the pre-dominant cell type present in the atherosclerotic plaque, and they constitute at least 30% of all human plaque cells^16^. Atherogenesis depletes SMC via suppression of SMC proliferation and increased cell death^16^. A pancake-shaped SMC phenotype was reported for type IV-VI lesions by AHA’s classification but not for normal vascular media^6^. We identified pancake-shaped αSMA^+^ cells in the tunica intima in each plaque group (Fig.2A) and the number of αSMA^+^ cells gradually decreased from 18.6±2.1% (group A) to 7.4±1.3% (group D) (Fig.2B).

EC dysfunction is important for development of atherosclerosis^17^ and EC depletion leads to formation of breaks in the endothelium contributing to plaque progression and destabilization. CAD also drives a process of new blood vessel development within atherosclerotic plaques called intraplaque neovascularization (IPN) that is linked to increased plaque vulnerability^18, 19^. To evaluate the relative level of breaks in the endothelial layer and quantify IPN, we immunostained coronary sections with antibody for CD31, endothelial cell marker^20^. We detected multiple single- and multi-cells breaks in CD31 signal in the luminal border in each plaque group. CD31-positivity in the luminal border area was reduced (P<0.05, ANOVA) in groups B, C and D compared to group A suggesting an increase in endothelial damage (Fig.2B). IPN was quantified in the tunica intima after subtraction of CD31-immunopositivity in the luminal border area. IPN was virtually undetected in group A (Suppl.Fig.4) and IPN signal was dramatically elevated in plaque group B to D (Fig.2B). We observed a significant >7-fold increase in IPN (P<0.001, t test) in plaque group C and D compared to group B lesions.

MF are a key player in progression of atherosclerotic lesions, regulating the local inflammatory milieu and plaque destabilization^21^. MF numbers increase multi-fold within mouse *aortae* during atherogenesis^22^. MF and MF-derived foam cells reached the largest number in the high*-*risk unstable plaques (type VI by AHA’s classification^6^). We identified SRA-positive cells in coronary plaques in the subendothelial layer and in the area surrounding the necrotic/lipid core (Fig.2B). The number of plaque SRA^+^ cells was gradually elevated in group A-D reaching the highest level in group D (15.1±1.7%). Plaques in group B-D have significantly higher SRA^+^ cells number compared to group A.

Collagen constitutes up to 60% of the total plaque protein thus contributing to plaque growth and the arterial lumen narrowing^23^. Collagen stimulates lesion progression by serving as a depot for pro-atherogenic molecules, by modulating macrophage functions, SMC proliferation and migration and by promoting thrombus formation^23^. On the other hand, a deficit of collagen leads to plaque weakness and vulnerability^24^. Collagen plaque level and location are an essential marker of fibrous plaque (fibroatheroma, type V by AHA’s classification)^5^. The fibroatheroma is defined as a lesion where a large necrotic/lipid core is encapsulated by a collagen-rich FC containing SMC and MF^5^. We detected high collagen levels in FC and around NC in porcine plaque group C and D (Fig.3A and see representative sections on Fig.1B). The collagen plaque level was the largest in group C (44.1±1.5%, Fig.3A).

**Figure 3.**
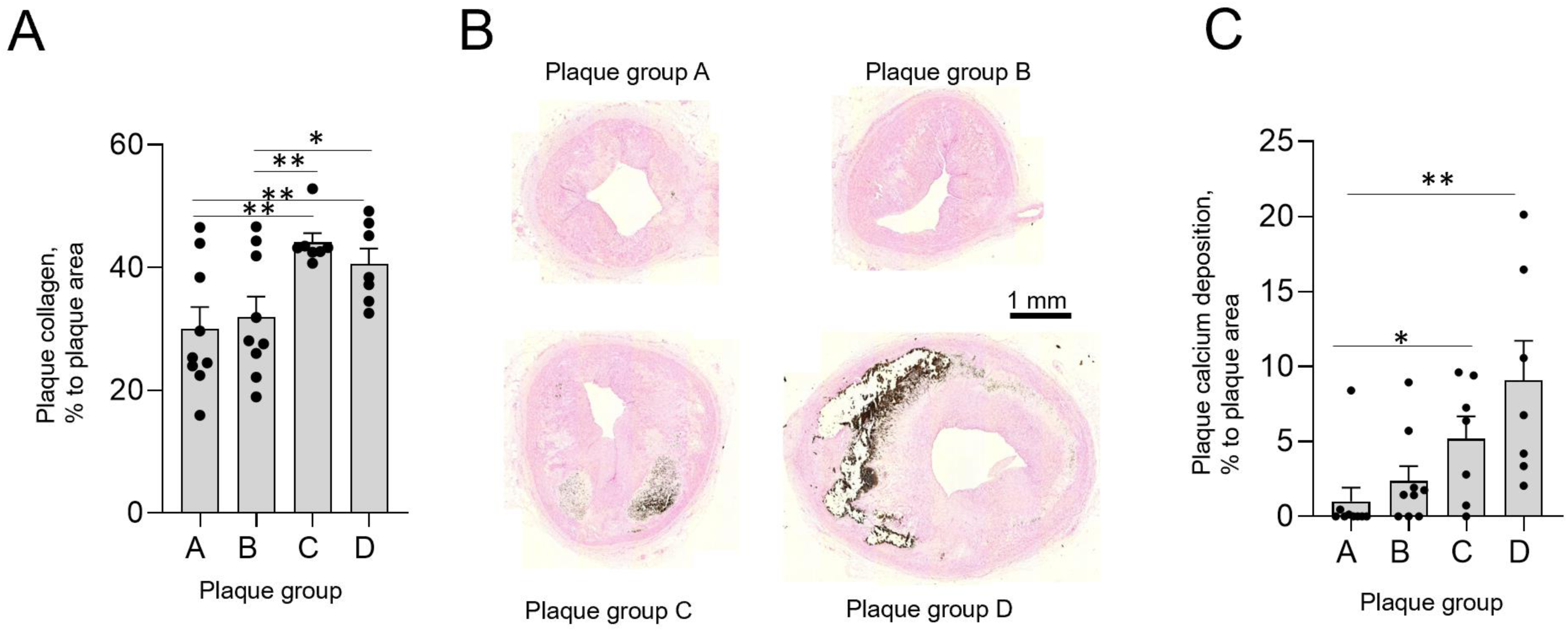
Collagen levels and calcium deposition in coronary plaques. Coronary sections (N=9/group A; N=9/group **8,** N=7/group C, N=7/group **D)** were stained with Trichrome or Von Kossa stain to detect collagen and calcium deposits, respectively. Collagen and calcium­ positive area were recognized by color-picking tool in CellSense software, and the area was normalized per tunica intima area. The collagen signal in plaque group **8-D** was normalized per plaque area after subtraction of necrotic core area. A, quantitative collagen data, **8,** representative images of Von Kassa-stained sections. C, quantitative data for calcium deposition. Scale bar, 1 mm. *P<0.05, **P<0.01, ***P<0.001, Student’s ttest.

It has been shown that ruptured human plaques had more calcifications compared to non-ruptured plaque^25^ suggesting the association of intraplaque calcification with advanced plaque phenotype and reduced plaque stability. However, there is no consensus regarding the specific role of calcium deposition for plaque stability. The current paradigm is that calcification of differential amounts, sizes, shapes, and positions may play differential roles in plaque homeostasis^26^. To test the potential association of calcium deposition with advanced plaque phenotype we performed quantification of calcium deposits in porcine coronary plaques. Plaques in group A and B have low levels of calcium deposition (<2%) in the form of diffuse single spots (Fig.3BC). Plaque calcification was increased multi-fold in plaque group C and D compared to group B (P<0.001, t test) suggesting the association of calcium deposition with advanced plaque phenotype. Calcified areas in human plaques are mostly located around the necrotic/lipid core close to the media^27^. We observed the most calcium deposits on the border of necrotic cores consistent with human studies.

### Features of vulnerable atherosclerotic plaque

The presence of well-formed fibrous cap, large necrotic/lipid cores and severe neovascularization are features of vulnerable plaque^28^. To obtain insights into plaque stability, all plaques were scored for the presence of fibrin-positive clots (thrombi) in intima, breaks in media or in fibrous cap, infiltrated large macrophage-shaped foam cells in the medial layer and RBC trapped inside tunica intima (Table 2 and Suppl.Fig.5). All plaques in group D and approximately 60% of plaques in group B and C contain abundant foam cells infiltrated in the tunica media. Importantly, 75% of plaques in group D also contain single or multiple thrombi inside intima. The number of plaques containing thrombi was reduced in group B (29%) and group C (20%). These results suggest that plaque group D represents an unstable plaque phenotype.

**Table 2.** Features of plaque vulnerability*.

| **Vulnerability index** | **A** | **B** | **C** | **D** |
| --- | --- | --- | --- | --- |
| Fibrous cap or media breaks | 0.08±0.05 | 0.21±0.11 | 0.19±0.11 | 0.31±0.13 |
| RBC trapped in intima | 0 | 0.07±0.07 | 0.13±0.09 | 0.16±0.10 |
| Single or multiple thrombi | 0 | 0.18±0.1 | 0.31±0.13 | 0.69±0.13 |
| Foam cells infiltrated in media | 0.33±0.08 | 0.54±0.13 | 0.69±0.13 | 1.00 |
| **Total score** | **0.42±0.13** | **1.00±0.41** | **1.31±0.46** | **2.16±0.36** |

\*Coronary sections were stained with Carstairs kit to detect fibrin, and plaque was scored for presence of breaks in fibrous cap or in media (+1), RBC trapped in tunica intima (+1), fibrin-positive thrombi in intima (+1), and/or foam cells infiltrated in media (+1) yielding the total score in range 0-4 for each plaque. The average and total score was calculated and shown in the table as mean±SEM.

### Porcine coronary plaque classification

We propose classification of porcine coronary lesions based on their structural features, histological and histochemical composition and features of plaque vulnerability (Table 3). Plaque group A does not contain FC or NC, has a relatively high level of SMC, and a low level of MF in the tunica intima. Plaque group A does not have a detectable level of neovascularization (Suppl.Fig.4). The plaque phenotype is consistent with the definition of intermediate (type III) human lesions^5^ that precede the development of advanced atherosclerotic lesions^29^. Type IV human lesions (atheroma) are the first plaque type considered advanced in AHA’s classification because of the presence of necrotic/lipid cores and severe intimal disorganization caused by the core formation^5^. Porcine plaque group B contains a relatively small (<15%) necrotic core and FC. In addition, we observed the presence of yellow-colored intracellular lipid droplets (foam cells) as well as extracellular lipid deposits in plaque group B, a feature of type IV human lesions. We conclude that plaque group B shares features with human type IV lesions.

**Table 3.**
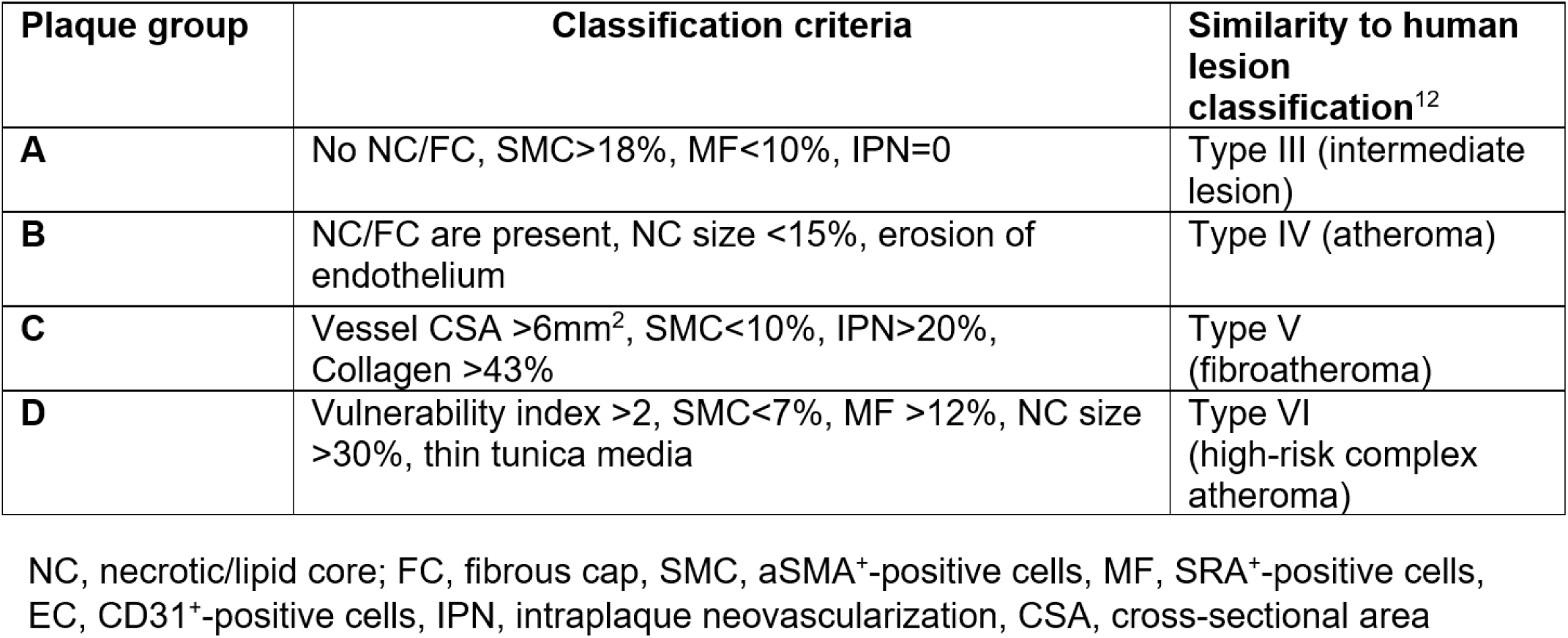
Porcine coronary lesion classification.

Human type V atheromas are defined as lesions in which new fibrous tissue has formed, and arteries are markedly narrowed with compensatory increase in total vessel size. The porcine plaque group C develops large necrotic/lipid cores (usually >2 cores), contains a thick (>0.5mm) fibrous cap, and significant level of neovascularization. The vessel has marked outward remodeling associated with thinning of the tunica media. Plaque C has the largest amount of Trichrome-positive collagen compared to other groups. These features of plaque group C are consistent with the definition of human type V lesions. Type VI human atheroma has all the features of type IV plaque plus developing breaks in the lesion or media, hematoma or hemorrhage, and thrombotic deposits^5^. The increase in plaque vulnerability in type VI atheroma is associated with depletion of SMC, increase in MF and increase in NC size^5^. Thus, the phenotype of porcine plaque group D is consistent with high-risk vulnerable type VI human atheroma. In summary, we assigned porcine group A, B, C and D to human lesion type III-VI, respectively (Table 3).

## DISCUSSION

The current report is the first to our knowledge to propose a classification of coronary atherosclerotic plaque in an animal model of atherosclerosis. To set up the algorithm for coronary plaque classification, we assessed changes in major morphological indices of atherosclerotic coronary arteries, i.e. IMR, vessel and plaque’s necrotic core size and fibrous cap thickness. Four plaque groups were identified by multi-featured clustering algorithm. Plaque groups were further characterized by analysis of cellular composition (SMC, MF and EC content), assessment of features of plaque vulnerability, quantification of lesion collagen, intraplaque neovascularization and calcification. Based on the similarity between porcine and human lesions, we assigned each porcine plaque group to a specific human lesion type. The reported classification aims to fulfill the following emergent needs in the field: 1) establish the suitability of the FH pig as a pre-clinical model mimicking advanced human CAD; 2) provide a clinically relevant research tool to assess changes in atherosclerotic lesion phenotype induced by interventions; 3) propose a practical framework for correlation of the composition of lesions with morphology determined by diagnostic imaging techniques.

We used intima-media ratio (IMR) as a discriminator of atherosclerotic plaque type. The increase in IMR correlated with elevation in intimal CSA showing the feasibility of using IMR as an *ex vivo* index of atherosclerotic burden. We found that atherosclerotic plaque progression induces thinning of the vascular media further supporting the usage of IMR for plaque grading. The medial thinning in coronaries was associated with infiltration of a large number of foam cells into the medial layer and appearance of multiple medial breaks, both signs of plaque destabilization. Our results are in line with reports showing loss of SMC and collagen in atherosclerotic tunica media^30^ and demonstration of attenuated contractile responses in a coronary artery with greater atherosclerosis^31^. Our result of an association of tunica media degradation with features of unstable plaque phenotype is a novel and intriguing finding. The classical concept of vulnerable plaque postulates that thinning of the fibrous cap is the main driving force of atheroma destabilization, thus potentially underestimating the role of the vascular media^1, 24, 28^. The thinning of the coronary vascular wall is known to promote the development of CAD-associated vascular pathologies such as coronary aneurysm (an abnormal dilatation of part of the coronary artery^32^) and coronary artery dissection^33^, however the mechanism underlying medial degradation and thinning is largely unknown.

Several laboratories have produced severe CAD in swine^34, 35^ demonstrating the feasibility of this model. Anti-atherosclerotic effects of novel drugs are routinely tested in pigs (including statins)^9, 36^. Unlike other animal models, pigs have human-like lipoprotein metabolism and swine also develop a coronary restenosis syndrome similar to humans^37^. Mild atherosclerotic lesions first appear in coronary arteries and both plaque distribution and composition follows a pattern comparable to that of humans^38^. Morphologic features of human advanced plaque including necrosis, calcification, neovascularization and intraplaque hemorrhage, were frequently reported for pigs^39, 40^ making pigs a valuable model to study plaque stability. Coronary atherosclerosis in FH swine occurs both spontaneously and by feeding with HFD for 4-6 months. We previously demonstrated that atherosclerotic plaque volume in porcine coronary arteries increased from 15% to ∼50-60% after 6 months of feeding with HFD and plaque composition at this time closely resembled human advanced atherosclerotic plaque^13^. Carotid plaques in FH pigs have high levels of expression of vascular cell adhesion molecule-1 and N-tyrosine (marker of oxidative stress)^41^ consistent with the vulnerable plaque phenotype found in humans^42^. Our current study provides important novel insights into coronary atherosclerosis development in FH pigs. We found that porcine plaque groups (A-D) follow a natural history of atherosclerosis established for human CAD. Our data suggests that transition between intermediate lesion and advanced atheroma in FH pig is characterized by change in vascular morphology closely mimicking human disease. In fact, our study identified the presence in pig coronary arteries of specific plaque phenotypes which were reported for humans. These findings promote the use of the FH pig as an experimental model developing the full spectrum of human-like coronary atherosclerosis.

Human histological fragments used for atherosclerotic plaque classification are often limited to autopsy samples collected from patients with end-stage CAD. These postmortem specimens have a relatively high prevalence of vulnerable plaques compared to samples obtained from animal models. Most human plaque samples are only partial vessel fragments limiting the accurate assessment of a total vessel’s tunica intima and media. Our assessment of coronary arteries in the FH pig allowed us to document atherosclerosis-induced changes in the vessel morphology (such as outward remodeling and media thinning) as well as analyzing true plaque heterogeneity over the length of the coronary artery. The current report provides an assessment of the full spectrum of atherosclerotic plaque types and disease-induced vascular remodeling which both are unavailable for most human histological studies.

Vulnerable plaques may rupture and thrombose, which is a life-threatening event, responsible for the development of the acute coronary syndromes of unstable angina, myocardial infarction, and sudden death^24^. The mechanisms underlying plaque destabilization includes increased cell death, defective efferocytosis (both promote the expansion of the necrotic/lipid core) and also neovascularization^43^. The role of cell senescence in plaque destabilization^44^, identification of “molecular signatures” of vulnerable plaque^45^ and determining the role of newly discovered cellular phenotypes (such as fibromyocytes and chondromyocytes^46^) are the focus of recent studies in the area of vulnerable plaque. Generation of a suitable animal model with a vulnerable plaque phenotype is a critical prerequisite to secure progress in the field. However, currently, there is no single, gold standard animal model of a vulnerable plaque^47^. FH pigs have great potential to advance vulnerable plaque research and provide quicker translation of results to patients. We report here that FH pigs on HFD have a significant number of group D lesions, which have multiple features of vulnerable plaque. We also found that plaque D has high IPN, MF levels, and prominent calcification, all features of high-risk atheroma. We had previously reported that some FH pigs had evidence of a myocardial infarction^13^ consistent with the presence of unstable coronary plaques. Novel revolutionary “omics” methods such as spatial transcriptomics (ST)^48^ open tremendous opportunities in the ongoing search for cell-specific determinants and “molecular signatures” of the thrombosis prone lesion and can boost progress in the generation of innovative anti-atherosclerotic therapies. In our previous report we used FH pigs as the first animal model of atherosclerosis to validate ST analysis with coronary plaque specimens^13^. ST confirmed the presence of recently identified fibromyocytes^49^ in the porcine coronary plaque. ST also detected histologically indistinguishable transcriptomic subclusters in the plaque’s fibrous cap. Taken together, these results show the great potential of the FH pig model in atherosclerosis-focused discovery research.

### Study limitations

First, coronary specimens used for the study were obtained only from female animals. Second, immunohistochemistry was used to assess plaque cellular composition and IHC is based on quantification of the relative expression of cell-specific markers. IHC does not allow accurate quantification of the number of marker-expressing cells. Third, the plaque classification misses the direct assessment of plaque lipids due to using paraffin-embedded cross-sections. Use of paraffin-embedded sections is an excellent choice for accurate vessel/plaque morphometry, however paraffin-embedded sections do not retain lipids, compromising direct lipid quantification. Fourth, the plaque vulnerability analysis includes the assessment of signs of former plaque rupture, however it is lacking data regarding whether the vessel was an infarct-related artery. Fifth, analysis of early lesions in FH pigs (similar to human type I and type II lesions) was not completed in the study since the animal protocol was optimized to promote development of advanced coronary atherosclerosis.

In summary, we have shown that FH pigs develop a full range of atherosclerotic plaques in coronaries and the plaque phenotype closely mimics human CAD. Our data support the use of the FH pig as a pre-clinical model with great potential to advance research in the field of CAD, especially in studying vulnerable plaque and in discovery research.

## ARTICLE INFORMATION

### AUTHORS CONTRIBUTION

SS designed the project and planned the experiments. ML performed plaque morphology assessment and clustering analysis. SS performed immunohistochemistry with coronary sections and plaque vulnerability assessment. SS wrote the original manuscript’s draft. PD supervised the project, handled funding, performed writing-review, and editing during manuscript preparation.

## ACKNOWLEDGEMENT

We thank to Dr. Douglas K. Bowles (University of Missouri at Columbia) and Dr. David Lefer (Cedars-Sinai Medical Center) for introducing us to FH pig model and support for all aspects of work with FH pigs. We acknowledge help of Dina Gaupp (Center for Gene Therapy at Tulane University Health Sciences Center) for making porcine sections and assistance with histological assays. We thank to all members of Dr. Delafontaine’s lab (Tulane University) for critical assessment of our findings.

