## Supplemental methods and figures for "Coronary atherosclerotic plaque composition and classification in hypercholesterimic pigs"

**SUPPLEMENTAL MATERIALS**

**METHODS**

Animals. *Rapacz* pigs with familial hypercholesterolemia (FH pigs) were used in our previous study to test the effect of insulin-like growth factor I (IGF-1) on atherosclerosis[1]. Fourteen-month-old FH pigs were administered with IGF-1 or saline (control) and fed with high-fat diet (HFD) for 6 months. Significant sex-specific differences in circulating lipids and in atherosclerosis development were found. FH pig females had higher cholesterol and triglyceride levels, and advanced coronary plaque phenotypes compared to FH males[1]. Right coronary artery (RCA) and left anterior descending artery (LAD) specimens collected from saline-injected female FH pigs (N=9) were used for the current study.

Coronary arteries morphometry. The proximal 30-35 mm fragment of RCA and LAD was fixed in 10% formalin, embedded in paraffin, and further cut onto six 5.0 mm fragments. A total of 108 vascular fragments (6 per coronary x 2 arteries x 9 pigs) were used to obtain cross-sections and stained with Trichrome Gomori's kit. Four fragments were discarded after initial inspection due to the presence of dissection-induced vascular damage, and a total of 104 fragments were submitted for analysis of vascular and plaque morphology. Slides were scanned with Zeiss Axio ScanZ.1 slide scanner. A slide with 1mm grid was included in each scanning session to assist with measurements. Analysis of vessel morphometry was performed with CellSens Dimension 1.18 software (Olympus). External elastic membrane (EEM), internal elastic membrane (IEM) and luminal border were manually outlined, and corresponding cross-sectional areas (CSA) were measured. The following indices were assessed:
Vessel CSA ($mm^2$) = EEM CSA-Luminal CSA,
Tunica media CSA ($mm^2$) = EEM CSA-IEM CSA,
Tunica intima CSA ($mm^2$) = IEM CSA-Luminal CSA,
The necrotic/lipid core (NC) area was manually outlined using CellSens software. The NC was defined as acellular (hematoxylin-negative) plaque area. The fibrous cap (FC) was defined as largely uninterrupted strip of brown-colored (smooth muscle, connective tissues) material on the top of the necrotic/lipid core with a higher density of nuclei than

the plaque necrotic/lipid core. The thickness of the FC was calculated as the mean length of 5 arbitrary lines distributed across the cap area. The intima-media thickness and intima-media ratio (IMR) were quantified by the following methods:

Method 1. The geometric center of outline for EEM, IEM and lumen was generated by CellSens software. The thickness of vascular media was calculated as a distance between EEM and IEM geometric centers, and intima thickness as a distance between IEM and lumen centers.

Method 2. The CSA of tunica intima (TICSA) and tunica media (TMCSA) were determined first. The radius (r) of circle corresponding to TICSA and TMCSA was calculated as r= √(TICSA/π) and r=√(TMCSA/π), respectively. The ratio of rTICSA/rTMCSA was calculated (IMR). IMR calculated by method 1 was in good agreement with one calculated by method 2 ($R^2$=0.94) (Suppl.Fig.1). We used IMR calculated by method 2 for plaque characterization in the current report.

Clustering analysis. The unsupervised K-means clustering multi-featured algorithm was used with R stats package v.2.15.3[2] to distinguish porcine coronary plaque groups. Four features of vessel/plaque morphology were selected for clustering: intima-media ratio (IMR), vessel area ($mm^2$), plaque's fibrous cap thickness (mm) and necrotic core area ($mm^2$). Prior to clustering, values of each feature were standardized using z-score normalization to ensure equal weighting in the analysis. To determine the optimal number of clusters we used two approaches: first, we applied the Elbow method[3] by calculating the Within-cluster Sum of Squares (WSS) criterion. The WSS measures the compactness of clusters by computing the total squared distances between data points and their assigned cluster centroids. The optimal number of clusters was identified at the point where adding more clusters yielded diminishing returns in reducing the WCSS. Second, we employed the Gap statistic method[4] to provide a statistical framework for cluster number selection. The Gap statistic compares the total within-cluster variation for different values of k with their expected values under a null reference distribution. We generated 50 Monte Carlo bootstrap samples to create the reference distribution. The optimal number of clusters was selected where the Gap statistic was maximized while accounting for statistical uncertainty through standard error estimation. Both Elbow and Gap statistical methods indicated that N=4 is the optimal number of clusters. Uniform Manifold Approximation and Projection (UMAP) algorithm was performed for normalized features to visualize clusters.

Immunohistochemistry (IHC) and histological assays. To perform IHC we selected sections for each plaque group with IMR in the 95% confidence interval range: N=9 sections/group A; N=9/group B, N=7/group C, and N=7/group D. Sections were incubated overnight at +4$^{o}$C with a mixture of primary antibodies, or with mixture of normal IgG (negative control). First, the section was incubated with mouse scavenger receptor A (SRA) antibody (TransGenia Inc., KT022, clone SRA-E5) plus rabbit CD31 antibody (Abcam, 134168, clone EP3095). SRA antibody signal was amplified with anti-mouse Alexa Fluor 647 Tyramide Super Boost kit (Invitrogen, B40916) and CD31 signal – with biotin-streptavidin system using anti-rabbit-biotin IgG (Abcam, BA-1000) followed by incubation with streptavidin-AlexaFluor 594 conjugate (Life Technologies, S32356) plus DAPI. Next, the section was incubated with α-smooth muscle actin antibody (αSMA) (clone 1A4)-AlexaFluor 488 conjugate (Invitrogen, 53-9760-82**)** and mounted with ProLong Gold antifade media (Thermo Fisher, P36970) for imaging. The fluorescent signals of AlexaFluor 647, AlexaFluor 594 and AlexaFluor 488 plus DAPI were quantified on the same section. We used Cytation 5 multi-mode imager (Bio-Tek, Winooski, MI) equipped with Cy5, Texas Red, GFP and DAPI filter cubes to generate greyscale images for each channel. The morphological regions-of-interest (ROI) (i.e., tunica media, necrotic/lipid core, etc.) were manually outlined and area immunopositive for each cellular marker was quantified within ROI. The ratio of marker-positive area per ROI area (x 100%) was calculated and shown in Figures.

Intraplaque neovascularization (IPN) was quantified using sections immunostained for CD31, an endothelial cell marker. First, the entire plaque and 50 um thick luminal border plaque area were manually outlined in CellSens software and CD31-immunopositive area was calculated for both luminal border and the entire plaque area. IPN was calculated as $CD31^{+}$-positivity per plaque after subtracting signal in the luminal border area. Trichrome staining is a quantitative method to detect muscle tissue (red) and collagen (blue)[5]. Plaque collagen was assessed using Trichrome-stained sections after outlining tunica intima area. Collagen-positive area was recognized by color-picking tool in CellSens software, and the area was normalized per intima area and shown as % in the graphs. Plaque calcification was assessed by staining of serial sections with Von Kossa stain[6] (Abcam, 150687) in according with manufacturer's instructions. Von Kossa detects calcium deposits as black spots on the pink background[6]. Intraplaque dark clusters were recognized by CellSens and the area was normalized per plaque area.

Assessment of features of vulnerable plaque. To quantify features of vulnerable plaque, coronary sections were stained with Carstairs kit (Electron Microscopy Sciences, 1965) in accordance with manufacturer's instructions. Carstairs stain detects fibrin, platelets, collagen, and red blood cells (RBC)[7]. Each plaque was scored for the presence of the following indices of plaque vulnerability: breaks in fibrous cap (FC) or in media (+1), RBC trapped in tunica intima (+1), fibrin-positive thrombi in intima (+1), foam cells infiltrated in media (+1) or none of them (0) yielding a total score in range 0-4 per plaque. The average vulnerability score per plaque and per group was calculated.

Statistical analysis. Statistical comparisons were performed by unpaired 2-tailed *t* test and using a repeated measures ANOVA. We specified the statistical test in each Figure legend. Grubb's *z* test was used to identify outliers. Data sets were first assessed for residual distribution using D'Agostino-Pearson omnibus normality test and for equal variances using Levene's test for equality of variances. Differences in outcomes were determined by ANOVA and Bonferroni's multiple comparisons test, Kruskal-Wallis test, or Mann-Whitney *U* test, according with the normality of residual distribution. For all comparisons, $P < 0.05$ was considered statistically significant. Data were analyzed using Microsoft Excel and Prism v.6.0 (GraphPad Software). Data presented in figures are individual data points (circles) and mean ± SEM (bars). Artwork was generated in GraphPad Prism and Adobe Photoshop 15.0.

1. Sukhanov S, Higashi Y, Yoshida T, Danchuk S, Alfortish M, Goodchild T, Scarborough A, Sharp T, Jenkins JS, Garcia D, et al. Insulin-like growth factor 1 reduces coronary atherosclerosis in pigs with familial hypercholesterolemia. *JCI Insight*. 2023;8. doi: 10.1172/jci.insight.165713
2. Team RC. *R: A language and environment for statistical computing. R Foundation for Statistical Computing*. Vienna, Austria; 2013.
3. Kuraria AJ, N.; Soni, M. Centroid Selection Process Using WCSS and Elbow Method for K-Mean Clustering Algorithm in Data Mining. *Int J Sci Res Sci Eng Technol*. 2018;4:190-195.
4. Tibshirani R. Estimating the Number of Clusters in a Data Set via the Gap Statistic. *Journal of the Royal Statistical Society Series B (Statistical Methodology)*. 2001;63:411-423.
5. Gurina TS, Simms L. Histology, Staining. In: *StatPearls*. Treasure Island (FL); 2025.
6. Schneider MR. Von Kossa and his staining technique. *Histochem Cell Biol*. 2021;156:523-526. doi: 10.1007/s00418-021-02051-3
7. Gu Y, Bai Y, Wu J, Hu L, Gao B. Establishment and characterization of an experimental model of coronary thrombotic microembolism in rats. *Am J Pathol*. 2010;177:1122-1130. doi: 10.2353/ajpath.2010.090889

**<u>Supplemental Figure 1.</u> Intima-media ratio (IMR) calculated by method 1 and 2.** IMR was quantified by method 1 as a ratio of geometric center of intima area per geometric center of media area or by method 2 as a ratio of radiuses calculated for circle corresponding intima and media area. IMR calculated by method 1 was in a good agreement with one calculated by method 2.

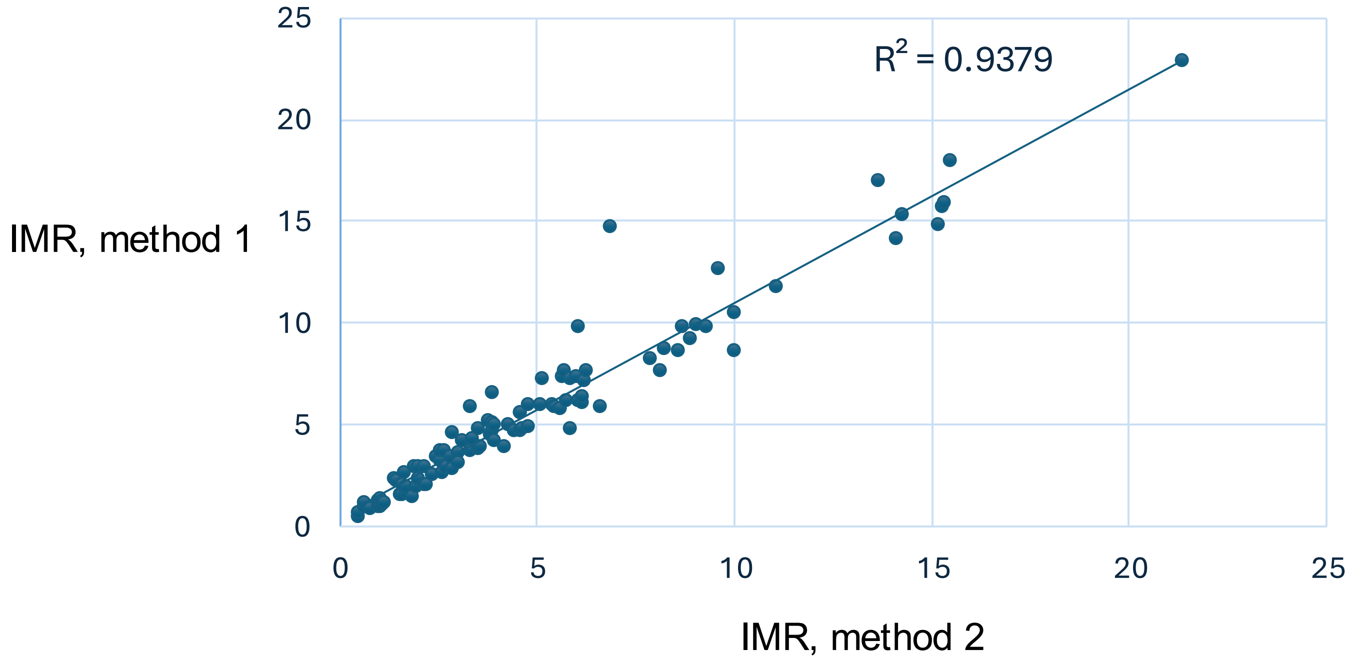

**<u>Supplemental Figure 2</u>. Determination of optimal number of clusters for multifeatured plaque clustering**. A, Elbow method calculates the Within-cluster Sum of Squares (WSS) criterion which is measures the compactness of clusters by computing the total squared distances between data points and their assigned cluster centroids. The optimal number of clusters was identified at the point where adding more clusters yielded diminishing returns in reducing the WCSS. B, Gap statistic method provides a statistical framework for cluster number selection. The Gap statistic compares the total within-cluster variation for different values of k with their expected values under a null reference distribution. The optimal number of clusters was selected where the Gap statistic was maximized while accounting for statistical uncertainty through standard error estimation. Both Elbow and Gap statistic method indicate that N=4 is the optimal number of clusters (arrow).

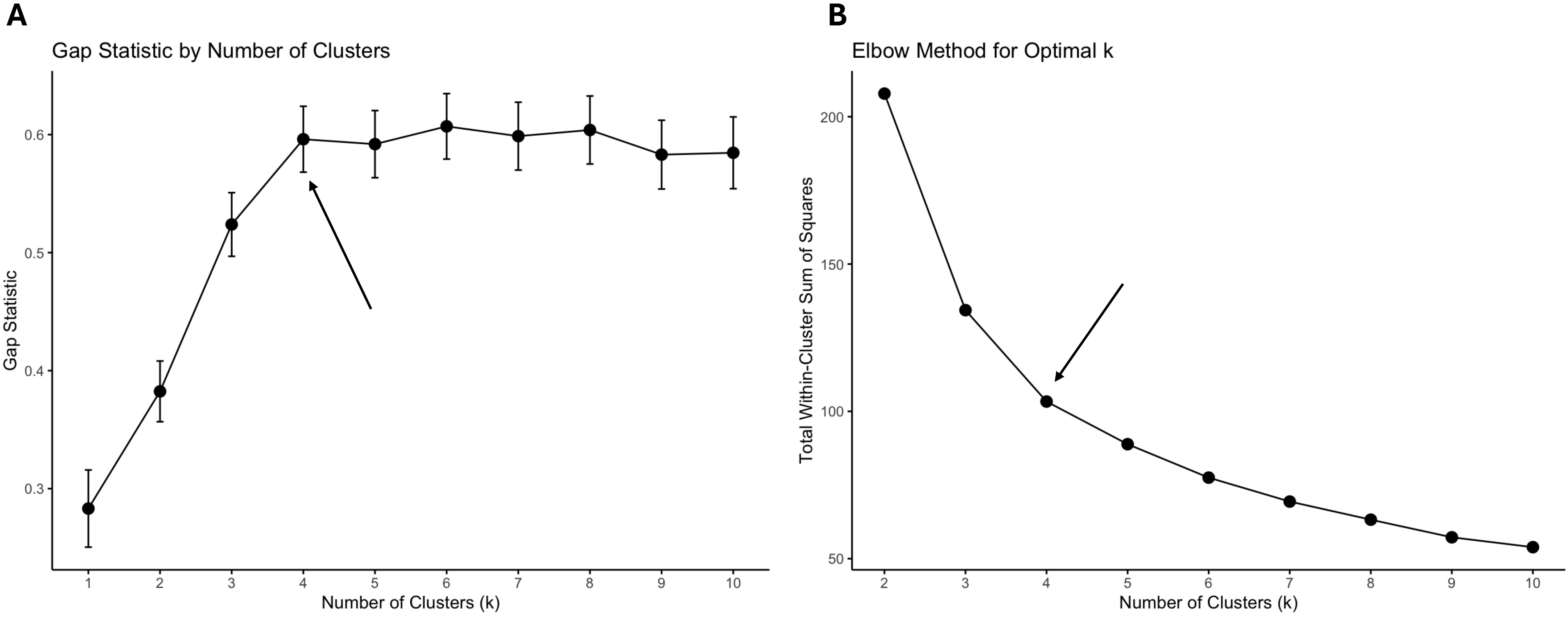

**<u>Supplemental Figure 3.</u> Porcine plaque clustering visualization using UMAP.**

Uniform Manifold Approximation and Projection (UMAP) algorithm was done with R kit for 4 normalized features to visualize clusters.

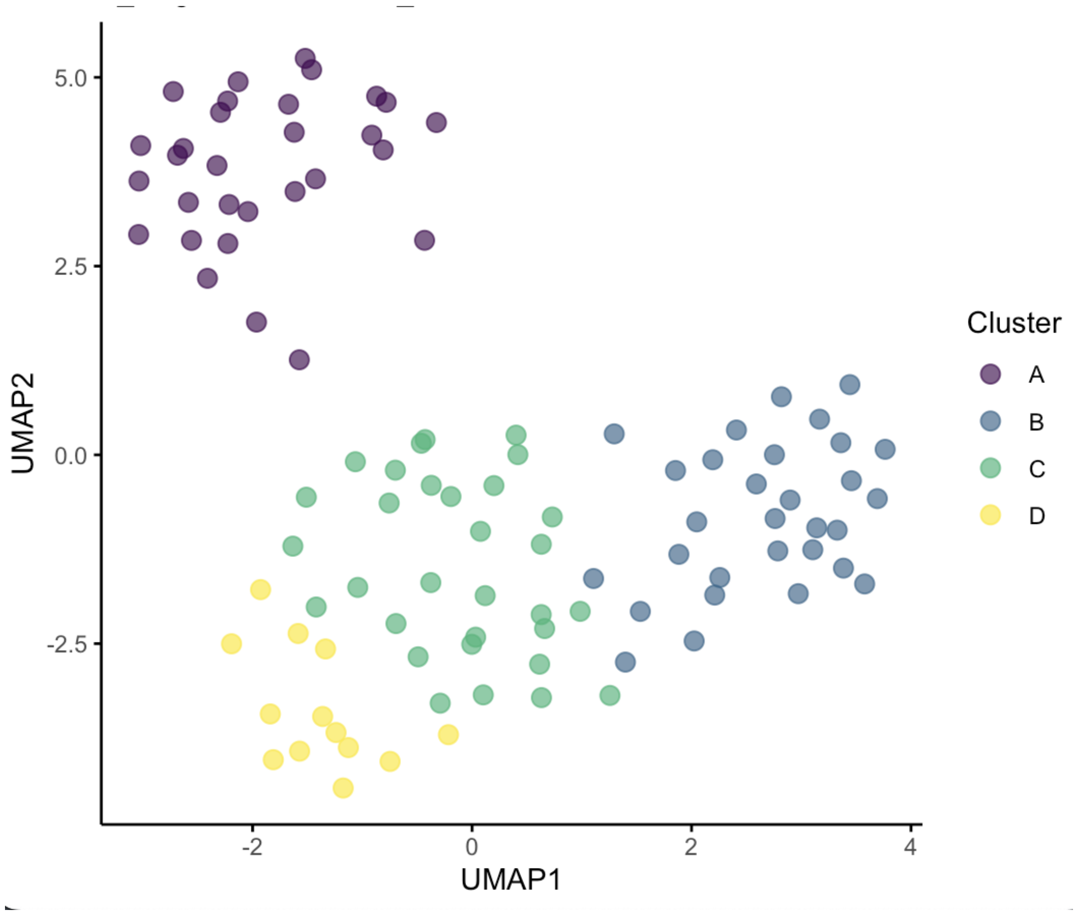

**Supplemental Figure 4. Intraplaque neovascularization (IPN).** Coronary sections were incubated with mixture of rabbit CD31 antibody (endothelial cell marker), α-smooth muscle actin antibody-AlexaFluor488 conjugate (α-SMA, smooth muscle cell marker) and DAPI. The CD31 signal was amplified with biotin-streptavidin system using anti-rabbit-biotin IgG and streptavidin-AlexaFluor 594 conjugate. IPN was detected in plaque group D and not in group A as a cluster of CD31+ cells inside tunica intima (yellow arrows). Scale bar, 1mm.

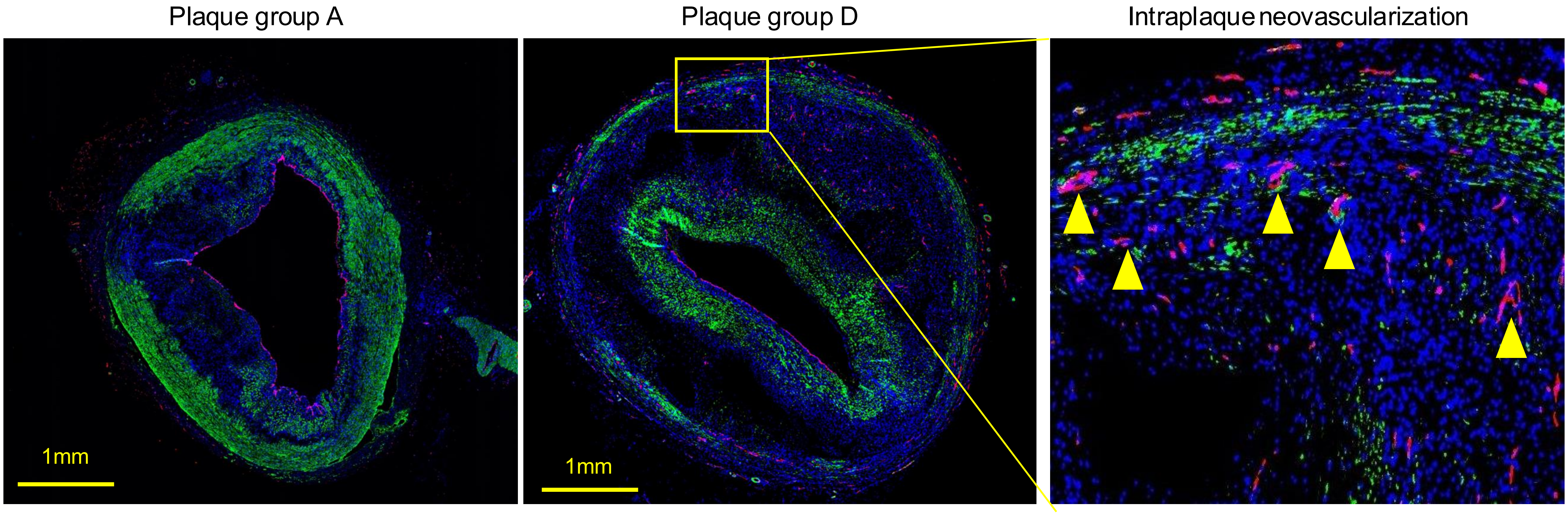

**Supplemental Figure 5. Features of vulnerable plaque**. Coronary sections were stained with Carstars stain to identify fibrin, platelets, collagen, and red blood cells (RBC). The presence of fibrin-positive thrombi (A), RBC trapped in tunica intima (B), foam cells infiltrated in media (C) and breaks in fibrous cap or in media was noted and scored. White arrows, thrombi; blue arrows, macrophage-shaped foam cells in vascular media; yellow arrows, break in tunica media (TM). Scale bar, 100 um.

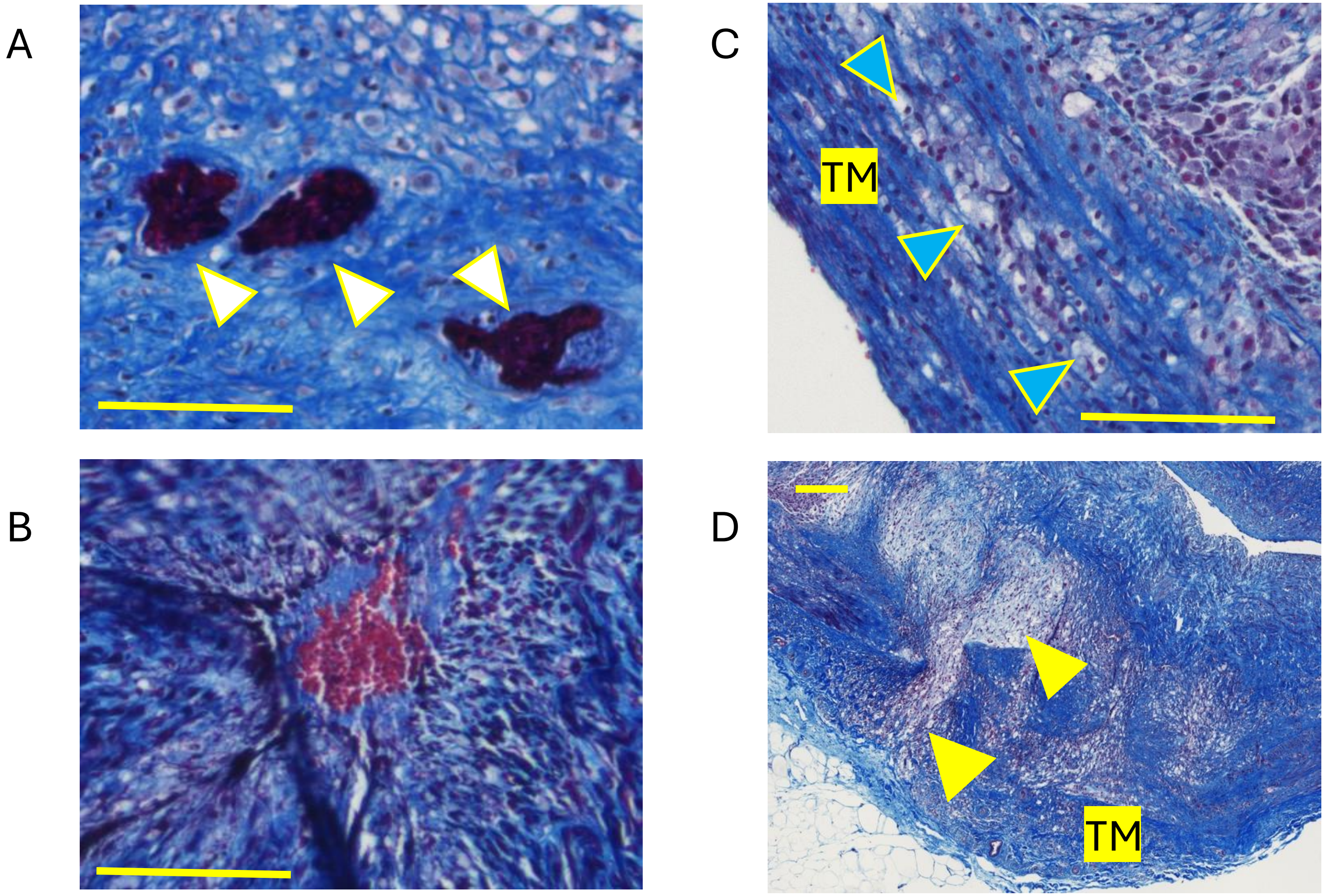
